# Drosophila Keap1 oxidative/xenobiotic response factor interacts with B-type lamin to regulate nuclear lamina and heterochromatin

**DOI:** 10.1101/2022.04.27.489742

**Authors:** Jennifer Carlson, Emma Neidviecky, Isabel Cook, Huai Deng

## Abstract

The essential function of the Keap1-Nrf2 pathway in mediating transcriptional response to xenobiotic and oxidative stimuli has been well established. However, the mechanisms whereby Keap1 and Nrf2 regulate developmental genes remains unclear. We hypothesized that *Drosophila* Keap1 (dKeap1) and Nrf2 (CncC) proteins regulate transcription through controlling high-order chromatin structure. Here, we describe evidence supporting that dKeap1 can regulate chromatin through interaction with lamin, the intermediate filament proteins that form nuclear lamina and organize the overall chromatin architecture. dKeap1 and lamin Dm0, the B-type lamin in *Drosophila*, interact with each other and form complexes in the nucleus. Overexpression of dKeap1 resulted in a redistribution of lamin Dm0 to the intra-nuclear area and consistently, caused a spreading of the heterochromatin marker H3K9me2 from the pericentromeric region to chromosome arms. Overexpression of dKeap1 fusion proteins in the *dKeap1* null background significantly disrupted the nuclear lamina morphology, indicating that dKeap1 is required for the maintenance of a normal nuclear lamina. Knock down of *dKeap1* partially rescued the lethality caused by lamin Dm0 overexpression, suggesting that dKeap1 and lamin Dm0 function in the same pathway during development. Taken together, these results support a model where dKeap1 regulates chromatin structure and developmental transcription through interaction with lamin proteins, revealing a novel epigenetic function of the Keap1 oxidative/xenobiotic response factor.

## Introduction

The Keap1-Nrf2 pathway plays an essential role in cell protection through mediating transcriptional responses to environmental toxins (xenobiotics) and oxidative stimuli (Itoh *et al*, 1999; Slocum & Kensler, 2011). Mis-regulation of Keap1 and Nrf2 can lead to a variety of diseases, including cancer (Taguchi & Yamamoto, 2017), respiratory diseases (Carlson *et al*, 2020), neurodegenerations (Uruno *et al*, 2020), and cardiovascular diseases (Smith *et al*, 2016). Nrf2 (NF-E2-Related Factor 2) is a transcription factor that activates a series of antioxidant and detoxifying genes (Malhotra *et al*, 2010; Taguchi *et al*, 2011; Zhang, 2006). Under basal conditions, Keap1 (Kelch-like ECH-associated protein 1) binds to Nrf2 in the cytoplasm and targets Nrf2 for ubiquitination and degradation. Xenobiotic/oxidative compounds disrupt the Keap1-Nrf2 interaction, thereby releasing Nrf2 to enter the nucleus and activate target genes (Eggler *et al*, 2005; Itoh *et al*., 1999).

Recent studies have found that Keap1 and Nrf2 can also regulate normal developmental programs in several model systems (Deng, 2014; Pitoniak & Bohmann, 2015). Some of these developmental functions are mediated by reactive oxygen species (ROS). ROS can serve as signals for developmental programs such as stem cell proliferation (Bigarella *et al*, 2014; Owusu-Ansah & Banerjee, 2009) and neuronal development (Oswald *et al*, 2018). As the master regulator of ROS, Keap1 and Nrf2 were found to regulate mouse intestinal stem cell proliferation and differentiation (Zhou *et al*, 2021). In *Drosophila*, dKeap1 and CncC (cap-n-collar C) proteins (the homologs of Keap1 and Nrf2, respectively) regulate proliferation of intestinal stem cells through ROS signaling (Hochmuth *et al*, 2011).

Keap1 and Nrf2 can also directly regulate transcription of developmental genes. For example, mouse Nrf2 and *Drosophila* dKeap1 both regulate adipogenesis through controlling adipogenesis genes (Carlson *et al*, 2022; Huang *et al*, 2010; Kim *et al*, 2018; Pi *et al*, 2010). Mouse Nrf2 also promotes cell proliferation by transcriptional activation of glucose metabolic enzymes (Mitsuishi *et al*, 2012) and promotes neuronal stem cell differentiation by activating genes that inhibit self-renewal (Khacho *et al*, 2016). CncC and dKeap1 promote *Drosophila* metamorphosis by activating ecdysone biosynthetic genes and response genes in specific tissues (Deng & Kerppola, 2013). The dKeap1-CncC signaling also plays a role in neuronal remodeling as CncC regulates dendrite pruning by activating gene expression of proteasomal subunits (Chew *et al*, 2021).

The mechanisms whereby Keap1/Nrf2 family proteins regulate developmental genes remain unclear. It is notable that in *Drosophila*, dKeap1 on one hand suppresses the CncC-activated transcription of detoxifying genes while on the other hand cooperates with CncC to activate some developmental transcripts (Deng & Kerppola, 2014). This indicates that the dKeap1-CncC complex employs distinct molecular mechanisms to control detoxification and development. Our previous studies have found that dKeap1 and CncC function at both euchromatin and heterochromatin regions. dKeap1 and CncC activate ecdysone-response genes through binding to the highly-decondensed “puffs” on the polytene chromosome (Deng & Kerppola, 2013). Both dKeap1 and CncC are also required for gene silencing at the pericentromeric heterochromatin (Carlson *et al*, 2019). Given that the roles of epigenetic mechanisms in the transcriptional regulation of development have been established (Kiefer, 2007), Keap1-Nrf2 family proteins could regulate developmental genes through epigenetic regulation of chromatin structure.

The nuclear lamina plays an important role in the organization and regulation of chromatin, especially heterochromatin. In general, euchromatic regions of the chromatin tend to be centrally localized in the nucleoplasm while heterochromatic regions tend to be associated with the nuclear lamina (Foster & Bridger, 2005). The nuclear lamina is assembled by lamin intermediate filament proteins. In mammals and *Drosophila*, there are two types of lamin proteins: A-type lamins (lamin A/C in mammals; Lamin C in *Drosophila*) and B-type lamins (lamin B1/B2 in mammals; lamin Dm0 in *Drosophila*) (Adam & Goldman, 2012). When all three mammalian lamins are knocked out in mice, euchromatin is relocated to the periphery of the nucleus and heterochromatin is centralized; this defect is partially restored by adding back lamin B1 (Zheng *et al*, 2018). Lamins can interact with chromatin either directly or indirectly (Hoskins *et al*, 2021; Shevelyov & Ulianov, 2019). For example, the tail of lamin Dm0 binds to histones H2A and H2B of nucleosomes (Goldberg *et al*, 1999). Lamin proteins can also regulate chromatin through interactions with other proteins such as HP1, LBR, Emerin, BAF, MAN1, LAP2β, PRR14, and NET (Shevelyov & Ulianov, 2019; Towbin *et al*, 2013).

The roles of lamin proteins in the transcriptional regulation of development have been established in different model systems. In mice, mutations in *lamin B1* result in abnormal lung development, defects in bone ossification, impaired differentiation of fibroblasts, and reduced viability (Vergnes *et al*, 2004). In humans, mutations in *lamin A* are associated with the Hutchinson-Gilford progeria syndrome, a genetic disease characterized by premature aging (Eriksson *et al*, 2003). In *Drosophila*, lamin Dm0 is required for male germline stem cell differentiation and proliferation (Chen *et al*, 2013). Mutations in *lamin Dm0* result in developmental defects in the ovary, central nerve system, and ventriculus (Osouda *et al*, 2005).

The developmental roles of lamins could be mediated by redox signals such as ROS, as several studies suggest that lamin proteins can both regulate and be regulated by ROS (Shimi & Goldman, 2014). For example, lamin B1 can regulate ROS levels through interaction with Oct-1, a transcription suppressor that silences antioxidant genes (Malhas *et al*, 2009). Lamins A/C and B1 reduce oxidative stress through the p53 pathway, thereby controlling cell proliferation of human diploid fibroblasts (Moiseeva *et al*, 2011; Shimi *et al*, 2011). On the other hand, oxidative stress activates the p53 pathway, which in turn, increases lamin A/C protein levels and reduces the level of lamin B1 (Shimi *et al*., 2011; Shimi & Goldman, 2014). A-type lamins also regulate levels of ROS. In *Drosophila*, mutations in *Lamin C* lead to cytoplasmic protein aggregation in muscle cells, which in turn results in reductive stress through the activation of CncC (Coombs *et al*, 2021).

Since Keap1 and lamin proteins both regulate heterochromatin, development, and oxidative responses, we investigated the hypothesis that Keap1 and lamin can co-regulate chromatin structure and developmental transcription. Here, we demonstrated a molecular and genetic interaction between dKeap1 and lamin Dm0 in *Drosophila*. We also found that dKeap1 overexpression re-localized lamin Dm0 into the intra-nuclear area and resulted in the spreading of the heterochromatin marker H3K9me2. Therefore, dKeap1 likely regulates heterochromatin structure and gene expression through its interaction with the nuclear lamina.

## Results and Discussion

### Molecular interaction between dKeap1 and Lamin

To explore the potential molecular function of dKeap1 in heterochromatin architecture, we tested whether dKeap1 associates with lamin Dm0 (also named as “Lamin” in Flybase, stated as “Lamin” in this article). Lamin Dm0 is the major type of lamin protein that forms the nuclear lamina in most of the *Drosophila* tissues. We first visualized the subcellular localization of dKeap1 and Lamin in salivary gland cells through co-immunostaining (Fig 1A). Although most of the dKeap1 immunosignals were in the nucleoplasm, co-localization of dKeap1 and Lamin signals were detected at the nuclear lamina, indicating that some dKeap1 proteins localize to the nuclear lamina.

**Figure 1.**
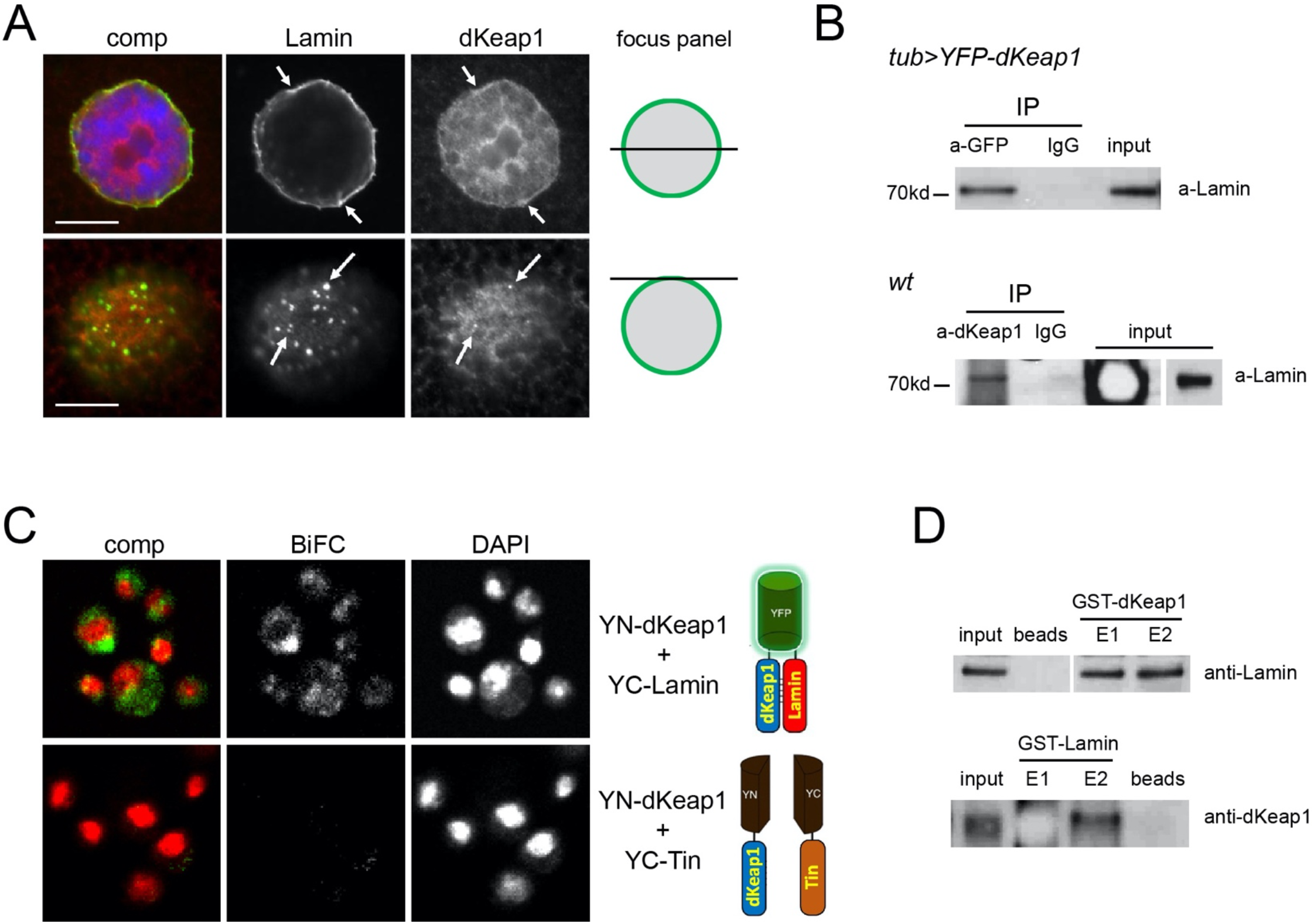
dKeap1 and Lamin interact and form a complex. *A. Subcellular localization of dKeap1 and Lamin*. Wildtype (Oregon R) salivary gland cells were immunostained with anti-lamin Dm0 (green) and anti-dKeap1 (red). DNA was stained with Hoechst (blue). The same nucleus was visualized in cross-section (top) and top surface (bottom). Selective loci where dKeap1 colocalized with Lamin are indicated by arrows. Scale bars: 10 μm. *B. Coimmunoprecipitation of dKeap1 and Lamin*. YFP-*dKeap1* fusion proteins expressed by *tub-GAL4* in embryos (upper) or endogenous dKeap1 proteins in wildtype embryos (lower) were precipitated with anti-GFP, anti-dKeap1 serum or IgG control. Lamin proteins were detected in the input and precipitations by western blotting. A lower exposure of the Lamin signal in the wildtype input was shown at right. *C. dKeap1-Lamin BiFC Complexes*. S2 cells were transfected with *YN-dKeap1* and *YC-Lamin* or *YN-dKeap1* and *YC-Tin* (negative control). YFP florescence (green) represents the formation of the dKeap1-Lamin BiFC complex. DNA was labeled by DAPI and pseudo-colored as red. The diagrams on the right depict BiFC analysis. *D. GST pull down of dKeap1 and Lamin*. GST-dKeap1 or GST-Lamin proteins were mixed with Lamin or dKeap1 proteins, respectively, and pulled down using Glutathione Sepharose 4B beads. As negative controls, Lamin or dKeap1 proteins were pulled down with beads only. The primary (E1) or secondary (E2) elutes from the beads were probed with anti-dKeap1 or anti-Lamin via western blotting.

We tested whether dKeap1 and Lamin are in the same protein complex using the co-immunoprecipitation assay. Lamin immunoblotting signal was detected in the anti-GFP precipitation from the lysate of embryos that expressed *YFP-dKeap1* (Fig 1B), indicating that the ectopic dKeap1 fusion interacts with Lamin. Anti-dKeap1 precipitation also co-precipitated Lamin proteins from the lysate of wildtype embryos (Fig 1B), suggesting that endogenous dKeap1 and Lamin form protein complexes *in vivo*.

We next visualized the formation of dKeap1-Lamin complexes in *Drosophila* S2 cells using the bimolecular fluorescence complementation (BiFC) assay (Hu *et al*, 2002). In the BiFC assay, the N-terminus and C-terminus of YFP (YN and YC) were fused to dKeap1 and Lamin, respectively (Fig 1C). Fluorescence signals representing dKeap1-Lamin BiFC complexes were detected in nuclei of S2 cells that co-expressed YN-dKeap1 and YC-Lamin (Fig 1C). As a negative control, no BiFC signal was detected in S2 cells that expressed BiFC fusions for dKeap1 and a cardiac transcription factor Tinman (Liu *et al*, 2009). Therefore, we concluded that dKeap1 and Lamin proteins can form specific complexes in the nucleus.

To examine whether dKeap1 and Lamin directly interact with each other, the *in vitro* GST-pull down assay was conducted (Fig 1D). GST-dKeap1 and GST-Lamin were expressed in and purified from *E. coli*. dKeap1 and Lamin proteins were generated by removal of the GST tag using protease. Both GST-dKeap1 and GST-Lamin were able to pull down Lamin and dKeap1, respectively (Fig 1D). Therefore, dKeap1 and Lamin physically interact with each other.

### Mis-regulated dKeap1 re-distributes Lamin and disrupts nuclear lamina

dKeap1-Lamin BiFC complexes were detected mainly in the nucleoplasm of S2 cells (Fig 1C). This is in contrast with the peripheral localization of endogenous Lamin in the nucleus (Fig 1A). We hypothesized that the mis-localization of dKeap1-Lamin BiFC complex was caused by the overexpression of dKeap1 fusion proteins. To test this possibility, we overexpressed the UAS-controlled dKeap1 full length (FL) fusion protein (Fig 1A) in salivary gland cells using *Sgs3-GAL4* (Cherbas *et al*, 2003). Significant amounts of Lamin proteins were detected in the nucleoplasm, with a pattern partially overlapping with dKeap1 fusion proteins (Fig 2B). The levels of Lamin proteins were not altered upon dKeap1 overexpression (Fig 2C). Therefore, overexpression of dKeap1 proteins caused a redistribution of Lamin proteins from the original peripheral sites to the center area of the nucleus.

**Figure 2.**
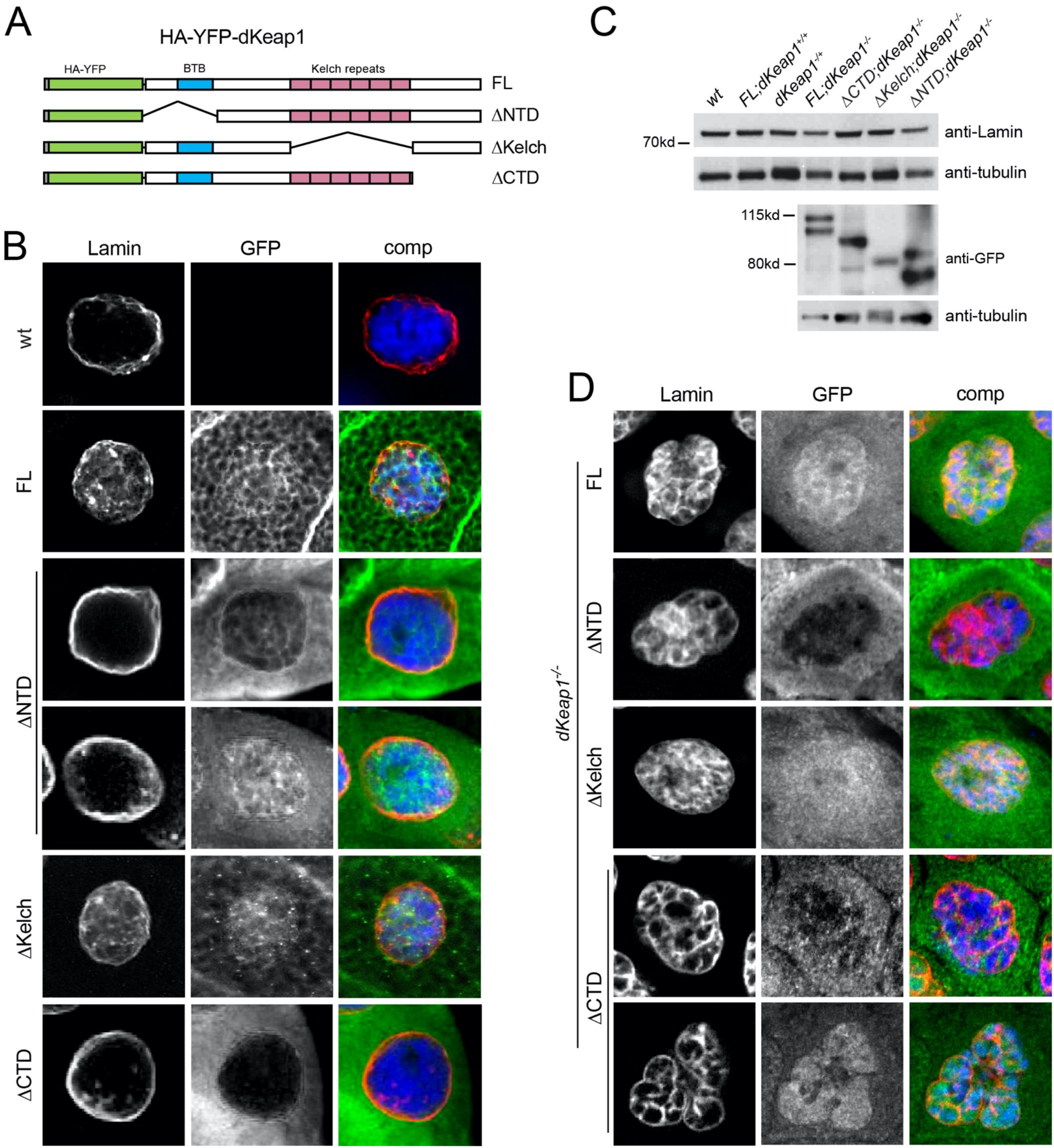
Mis-regulated dKeap1 relocalizes Lamin and disrupts nuclear lamina. *A. Diagram of dKeap1 fusion proteins*. dKeap1 full length (FL) and truncated proteins including ΔNTD (missing the N-terminal and BTB domain), ΔKelch (missing the Kelch repeats), and ΔCTD (missing the C-terminal domain) were tagged by HA and YFP. *B. Effects of dKeap1 overexpression on Lamin in wildtype background*. UAS-controlled dKeap1 fusion proteins described in A were expressed in the salivary glands via *Sgs3-GAL4*. Resulting salivary glands were immunostained with anti-GFP (green) and anti-Lamin (red). DNA was stained with Hoechst (blue). *C. Lamin protein levels in dKeap1 mutant flies*. Protein levels of Lamin and dKeap1 fusions in L3 larvae with genotypes described above were examined via western blotting using anti-Lamin and anti-GFP, respectively. Anti-tubulin was used as a loading control. *D. Effects of dKeap1 overexpression on Lamin in a dKeap1 null background*. UAS-controlled dKeap1 fusion proteins described in A were expressed in the *dKeap1*^*EY5/EY5*^ null mutant using *tub-GAL4*. Resulting salivary glands were immunostained with anti-GFP (green) and anti-Lamin (red). DNA was stained with Hoechst (blue). *EV2A. Mis-regulated dKeap1 disrupts Lamin organization in ovary cells. Ovaries from wildtype flies or flies expressing YFP-dKeap1* or *YFP-dKeap1-*Δ*CTD* in the *dKeap1*^*EY5/EY5*^ null background using *tub-GAL4* were immunostained with anti-Lamin. In both diploid follicle cells and polyploid nurse cells, nuclear lamina morphologies were abnormal and intra-nuclear localizations of Lamin proteins were detected. *EV2B. dKeap1 knockout has no effect on Lamin morphology in early larvae. Tissues from early L1 larvae of dKeap1*^*EY5/EY5*^ null mutants were immunostained with anti-Lamin (red). DNA was stained with DAPI (blue).

A similar Lamin redistribution phenotype was also observed in salivary gland cells that overexpressed YFP-dKeap1-ΔKelch, a dKeap1 truncation lacking the CncC-interacting Kelch domain (Figs 2A, 2B). The signals of YFP-dKeap1-ΔKelch were mainly detected in the nucleus. As this truncation induced the same effects as YFP-dKeap1-FL did, we concluded that the dKeap1-CncC interaction is not involved in the regulation of lamina by ectopic dKeap1. Expressing fusions of dKeap1 N-terminal deletion (YFP-dKeap1-ΔNTD) or C-terminal deletion (YFP-dKeap1-ΔCTD) had no or only moderate effect on Lamin distribution. YFP-dKeap1-ΔCTD localized almost exclusively to the cytoplasm (Fig 2B) (Carlson *et al*., 2022). YFP-dKeap1-ΔNTD localized to both the cytoplasm and nucleus, and the ratio of different portions varied in different cells (Fig 2B). The nuclear accumulation of dKeap1-ΔNTD had no significant effect to the Lamin distribution, indicating that the N-terminal domain of dKeap1 is required for the relocation of Lamin. The expression levels of all the fusion proteins were comparable and none of them altered Lamin protein levels (Fig 2C), indicating that ectopic dKeap1 proteins induced Lamin re-distribution directly rather than altering Lamin protein levels.

Severely defected nuclear lamina morphologies were observed when these dKeap1 fusion proteins were expressed in a *dKeap1* null background (Fig 2D). Interestingly, all the truncations, when expressed in the *dKeap1* null background, caused severe Lamin redistribution to the nucleoplasm (Fig 2D). In addition, expression of dKeap1-FL, ΔNTD or ΔCTD also altered the morphologies of nuclear laminas and the shapes of nuclei (Fig 2D), regardless of whether the fusion proteins were primarily in the nucleus or cytoplasm. In some cells that expressed ΔCTD, nuclear shapes and organizations were the most severely affected (Fig 2D). These cells showed partial nuclear fragmentation and potential breakdown of the nuclear envelope as indicated by the invading of ΔCTD fusion proteins from the cytoplasm into the nucleus (Figs 2B, 2D). The re-distributed Lamin and defected nuclear lamina were also seen in other cell types such as the diploid follicle cells and polyploid nurse cells in ovaries (Fig EV2A).

The lack of native dKeap1 proteins in the *dKeap1* null background should account for the severe Lamin defects induced by ectopic dKeap1 fusions. However, no significant Lamin defect was detected in cells of the *dKeap1* null larvae (Fig EV2B). Intrinsic dKeap1 proteins localized to both the nucleoplasm and the nuclear lamina, while none of the dKeap1 fusion proteins showed localization to the nuclear lamina (Fig 1A, 2B), indicating that the YFP-dKeap1 fusion proteins cannot function the same as intrinsic dKeap1 proteins in the nuclear lamina. In support of this, overexpression of YFP-dKeap1-FL largely but cannot fully rescue the viability and fertility of the *dKeap1* null mutant (Carlson *et al*., 2022). It is possible that the HA-YFP tag interferes with the interaction between dKeap1 and Lamin. Therefore, the disrupted Lamin morphology is likely a combinatory effect of both the ectopic expression dKeap1 fusion proteins and the lack of endogenous dKeap1 proteins. Since that the dKeap1-ΔCTD induced the worst nuclear lamina disruption, the C-terminal domains of dKeap1 should play the most significant role in the maintenance of a normal nuclear lamina shape.

Several other mis-localized lamin phenotypes have been reported. In *Drosophila*, reduction of a WAS family protein wash in salivary gland cells results in reduced nuclear envelope budding and patternless strands of Lamin throughout the nucleoplasm (Verboon *et al*, 2020). Overexpression of an inner nuclear membrane protein Kugelkern in *Drosophila* embryos and mouse fibroblasts results in infoldings of the nuclear membrane and “blebs” of the nuclear envelope, respectively (Polychronidou *et al*, 2010). Notably, expressing a lamin Dm0-Δ428-451 truncation (*nlsLamD*Δ*I*) in *Drosophila* salivary gland cells results in a defect described as invaginations and lobulations in the lamina (Uchino *et al*, 2017). This phenotype is similar but not identical to the dKeap1-induced defects seen in this study. It is possible that dKeap1 interacts with the chromatin binding region between the rod domain and the tail domain of Lamin. *Lamin A* mutations in human skin fibroblasts show nuclear lamina phenotypes described as “blebs”, “donuts”, and “honeycomb” at one end of the nucleus (van Tienen *et al*, 2019). *Lamin B1* mutations in mouse embryonic fibroblasts result in “honeycomb” throughout the whole nucleus (Jung *et al*, 2013). Compared to these “honeycomb” phenotypes, the dKeap1-induced honeycomb-like lamina defect revealed in our study shows much larger “holes” throughout the entire nucleus.

Intra-nuclear distributions of lamin proteins have been found in embryonic stem cells, presumably associated with a distinct global chromatin architecture in these pluripotent cells (Meshorer & Misteli, 2006). Hematopoietic stem cells from mouse adult bone marrow and human fetal bone marrow have misshapen nuclei with intra-nuclear lamin A/C (Dorland *et al*, 2019). Intra-nuclear lamins and abnormal nuclear lamina are also revealed in aging cells (Scaffidi & Misteli, 2006). For example, intra-nuclear lamin A proteins are found in normal human old fibroblasts and in fibroblasts from individuals with Hutchinson-Gilford progeria (Scaffidi & Misteli, 2006). Interestingly, conditional expression of Lamin or Kugelkern in *Drosophila* adult tissues reduces the lifespan and causes wrinkled and lobulated nuclear shapes (Brandt et al. 2008). Overexpression of dKeap1 fusion proteins allowed the early lethal *dKeap1* null mutants to survive to adults, but the rescue was only partial, and these flies showed reduced fertility (Carlson *et al*., 2022). We hypothesize that this developmental defect is associated with the disrupted lamina structure and the consequent mis-organization of chromatin and mis-regulation of developmental genes.

### dKeap1 overexpression redistributes heterochromatin marker H3K9me2

Our previous studies found that knockout of *dKeap1* reduced the level of heterochromatin marker histone H3K9me2, suggesting that dKeap1 is required for heterochromatin formation and/or maintenance (Carlson *et al*., 2019). Given that dKeap1 overexpression relocated Lamin, we tested whether ectopic dKeap1 can alter the organization of heterochromatin in the nucleus. We investigated the level and distribution of H3K9me2 in salivary gland cells that expressed YFP-dKeap1. In wildtype cells, H3K9me2 is found at the chromocenter, the pericentric heterochromatin region on the polytene chromosome (Fig 3A) (Zhang *et al*, 2006). Significant relocations of H3K9me2 immuno-signals to loci outside of the chromocenter were detected in nuclei with dKeap1 overexpression (Fig 3A, more examples shown in Fig EV3). Western botting assays showed that although *dKeap1* knockout reduced the level of H3K9me2, dKeap1 overexpression had no effect on the level of H3K9me2 (Fig 3B). Therefore, dKeap1 overexpression caused spreading of the H3K9me2 from the pericentric region to chromosome arms. We suggest that this relocation of H3K9me2 heterochromatin marker is a consequence of ectopic dKeap1 and Lamin accumulation in the center area of the nucleus. These results further support that the dKeap1-Lamin complex may coregulate transcription through controlling chromatin architerture.

**Figure 3.**
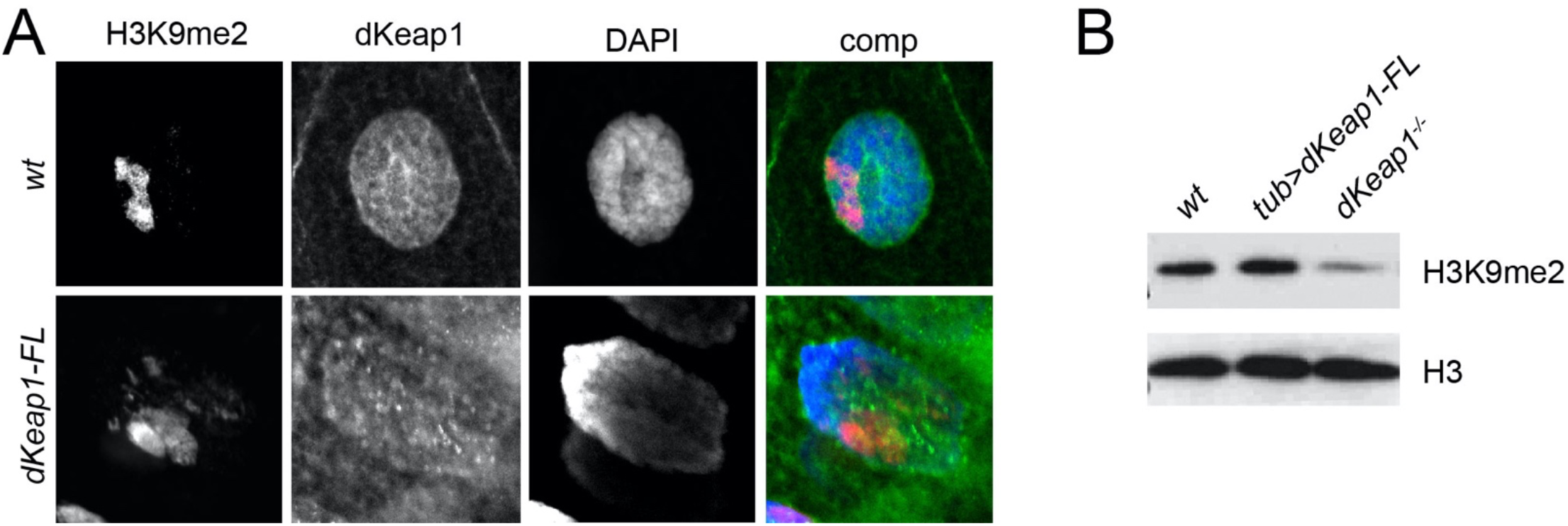
dKeap1 overexpression redistributes heterochromatin marker H3K9me2. *A. H3K9me2 distributions in salivary gland nuclei*. Salivary glands from wildtype larvae or larvae that expressed *YFP-dKeap1-FL* using *tub-GAL4* were immunostained with anti-H3K9me2 (Red) and anti-GFP (green). DNA was stained with DAPI. *B. Levels of H3K9me2 in dKeap1 overexpression or knock out larvae*. H3K9me2 levels were detected in early L1 larvae (the *dKeap1*^*EY5/EY5*^ null mutant is L1 lethal) with genotypes described above using western blotting. YFP-dKeap1-FL was overexpressed via *tub-GAL4*. Histone H3 was used as the loading control. *EV3. dKeap1 overexpression spreads heterochromatin marker H3K9me2*. Salivary glands from larvae that expressed *YFP-dKeap1-FL* via *tub-GAL4* in the wildtype background were immunostained with anti-H3K9me2.

### dKeap1 functions downstream of Lamin in the genetic pathway

To determine if dKeap1 and Lamin regulate gene expression in the same developmental pathway, we explored potential genetic interactions between *dKeap1* and *Lamin*. Lamin overexpression causes early lethality in *Drosophila* (Munoz-Alarcon *et al*, 2007). In our experiments, larvae that overexpress Lamin driven by *tub-GAL4* died at L1 or early L2 larval stage (Fig 4A). Double mutants which contained overexpressed Lamin and a heterozygous *dKeap1* null allele survived to late L2 or early L3 stage (Fig 4A), indicating that reduction of dKeap1 was able to partially rescue the lethality caused by Lamin overexpression. Excessive Lamin proteins could cause lethality via mis-regulation of developmental transcription. These results suggest that dKeap1 can act down-stream of Lamin in the regulation of gene expression during *Drosophila* development.

**Figure 4.**
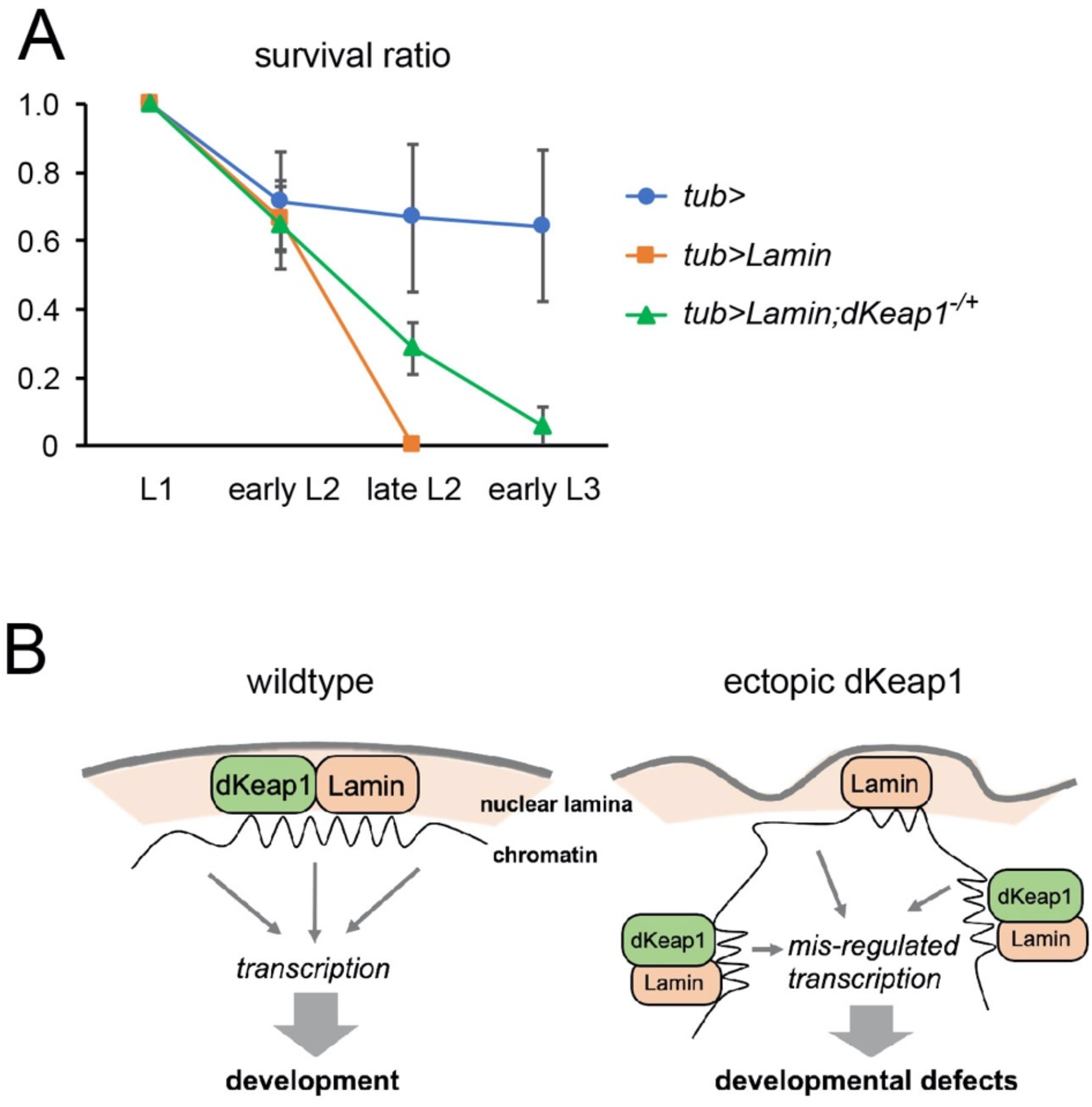
dKeap1 and Lamin function in the same genetic pathway. *A. Genetic interaction between dKeap1 and Lamin mutants*. Survival ratios of control flies (*tub-GAL4/+*), Lamin overexpression flies (U*AS-YFP-Lamin/+; tub-GAL4/+*), and the double mutant combining Lamin overexpression and dKeap1 knock down (*UAS-YFP-Lamin/+; dKeap1*^*EY5*^,*tub-GAL4/+*) were measured at larval stages listed below. Error bars represent the standard deviation based on two independent experiments. T-tests were used to compare the survival ratios of *Lamin* overexpression flies (*tub>Lamin*) and the double mutant flies (*tub>Lamin;dKeap1*^*-/+*^) (p<0.05). *B. Model of dKeap1-Lamin interaction and function*. The interaction between dKeap1 and Lamin proteins at the nuclear lamina could mediate the regulation of chromatin architecture, especially heterochromatin. This would indicate a potential epigenetic mechanism whereby Keap1 family proteins regulate developmental genes. Ectopic dKeap1 proteins relocate Lamin proteins to the intra-nuclear area, inducing mis-regulated transcription in development. dKeap1 is also required for the maintenance of normal nuclear lamina morphology.

Mutations in *lamin Dm0* show reduced viability, abnormal tissue differentiation, and defects in fertility, ovary size, ventriculus, and locomotion (Lenz-Bohme *et al*, 1997; Munoz-Alarcon *et al*., 2007; Osouda *et al*., 2005). A dKeap1 mutant with disrupted dKeap1 chromatin binding also shows reduced viability and fertility (Carlson *et al*., 2022). It would be interesting to determine the developmental genes and programs that are coregulated by dKeap1 and Lamin.

### Model of dKeap1-Lamin interaction and function

In contrast to the well-established model in which Keap1 and Nrf2 factors control antioxidant and detoxifying transcription, the mechanisms whereby Keap1 and Nrf2 regulate developmental genes remains unclear. Keap1 and Nrf2 likely function both in gene activation as transcription activators and in gene silencing by regulating heterochromatin (Carlson *et al*., 2022; Carlson *et al*., 2019; Deng & Kerppola, 2013, 2014). We hypothesize that Keap1-Nrf2 conducts different roles by forming complexes with different protein partners. In this study, we revealed that dKeap1 interacts with Lamin, and this novel interaction supports a model in which the dKeap1-Lamin complex controls the organization of heterochromatin and the overall chromatin architecture in the nucleus (Fig 5B). This finding would also indicate a novel epigenetic mechanism whereby Keap1 family proteins regulate developmental genes (Fig 5B). Our model is directly supported by several observations. First, dKeap1 and Lamin molecularly interact with each other and form complexes in the nucleus. Second, ectopic dKeap1 expression causes a relocation of Lamin proteins from the periphery regions to the center area of nuclei. dKeap1 is also required for the maintenance of normal nuclear lamina morphology. Third, ectopic dKeap1 expression causes a spreading of the heterochromatin marker from the pericentric heterochromatin region to the euchromatic polytene chromosome arms. Finally, the genetic interaction between *dKeap1* and *Lamin* suggests that they function in the same pathway during development.

Although it has been established that xenobiotic and oxidative stresses can affect development and lead to cancer and other diseases (Barouki *et al*, 2012), the underlying molecular mechanisms are not yet fully elucidated. It is thought that epigenetic mechanisms mediate the influence of environmental toxins on development at the transcriptional level. Recent environmental epigenetic studies have revealed broad influences of environmental toxins on human epigenomes and disease (Feil & Fraga, 2012; Marsit, 2015). However, the molecular factors that bridge the epigenetic alterations and environmental stimuli are less understood. Keap1 family proteins are the primary cellular sensors for oxidative and xenobiotic stimuli. Revealing the novel role of the dKeap1-Lamin complex in chromatin organization and gene regulation luminates a hypothesis in which the dKeap1-Lamin pathway mediates epigenetic and transcriptional adaptations to environmental toxins. It remains to be explored whether the dKeap1-Lamin complex responds to oxidative/xenobiotic stimuli, how dKeap1-Lamin controls nuclear chromatin architecture and the epigenome, and the full range biological functions of the dKeap1-Lamin pathway. These studies are also expected to broaden our understanding of the complicated roles of Keap1 and Nrf2 in human diseases.

## Materials and Methods

### Drosophila stocks

Fly stocks were maintained at 25°C according to standard protocol. Constructs of *UAS-YFP-dKeap1-FL, UAS*-*YFP-dKeap1-*Δ*NTD, UAS-YFP-dKeap1-*Δ*Kelch*, or *UAS-YFP-dKeap1-*Δ*CTD* were generated in pUAST vector and injected into the *w*^*1118*^ background. *Sgs3-GAL4, tub-GAL4*, and *UAS-YFP-Lamin* (BL7376) were from the Bloomington Stock Center. *dKeap1*^*EY5*^ was provided by Dirk Bohmann (Sykiotis & Bohmann, 2008). *dKeap1*^*EY5*^ and *tub-GAL4* are combined with the *TM6,Tb,Sb,Hu,e,Dfd-YFP (TM6)* balancer. Appropriate progenies were identified based on *Tb* marker in larvae and pupae or *Sb* marker in adults. Embryos with or without the *Dfd-YFP* marker were sorted under a Leica MZ10 F fluorescence stereomicroscope.

### Immunofluorescence

Salivary glands isolated from L3 larvae and ovaries isolated from adults are fixed in 3.7% paraformaldehyde for 5 minutes. Tissues were then washed with PBST (PBS + 0.2% Triton X-100) and stained with anti-lamin Dm0 (1:500, ADL67.10, Developmental Studies Hybridoma Bank) or anti-GFP (1:200, Novus, NB600) at 4°C overnight. After washing with PBST, the tissues were stained with 1:2000 goat anti-mouse Alexa Fluor 594 and goat anti-rabbit Alexa Fluor 488 secondary antibodies (Invitrogen) at 25°C for 2 hours, washed with PBST, and then stained with Hoechst in PBS for 10 minutes. Slides were mounted in Vectashield (Vector Laboratories) and imaged using a Zeiss LSM 710 confocal microscope.

### Co-immunoprecipitation

*UAS-YFP-dKeap1/+; tub-GAL4/+* embryos were collected on apple juice plates. Approximately 2000 embryos were ground in IP buffer (20 mM Tris-HCl pH 8.0, 150 mM NaCl, 11 mM EDTA, 0.2% Triton X-100, 0.2% NP-40, 2 mM Na_3_VO_4_, 1 mM PMSF, and 1.5 μg/ml aprotinin). 10 μl of GFP antibody (Novus, NB600) or negative control antibody normal rabbit IgG (Cell Signaling) was added to the cell lysate and the samples were rotated at 4°C overnight. For each IP, 30 μl Protein G agarose beads (Cell Signaling) were washed in IP buffer and blocked with bovine serum albumin (BSA), then added to the lysate-antibody solution and rotated for 2 hours at 4°C. Beads were spun down and the proteins were eluted using SDS-PAGE dye at 100°C for 10 minutes, followed by western blotting.

### Western blotting

In all western blots, 20 μl of sample were loaded in a 4-12% Bis-Tris gel (Invitrogen) and run for 2.5 hours at 110V. Proteins were transferred to nitrocellulose membranes (Bio-Rad) at 11V for 17 hours at 4°C. Membranes were blocked with 5% milk in TBST (TBS + 0.1% Tween-20) and then probed with primary antibodies against lamin Dm0 (1:400, Developmental Studies Hybridoma Bank ADL67.10), dKeap1 (1:100)(Deng & Kerppola, 2013), tubulin (1:500, Developmental Studies Hybridoma Bank 12G10), H3K9me2 (1:200, abcam ab1220), Histone H3 (1:1000, Proteintech 17168), GFP (1:500, Novus NB600), or GST (1:500, Genscript A00865), followed by 1:3000 HRP-conjugated Goat anti-rabbit or Goat anti-mouse secondary antibodies (Bio Rad). The membranes were developed by ECL reagents (GE Healthcare) and then exposed to X-ray film (AGFA).

### GST pull down

Full length *dKeap1* and *Lamin* were cloned into pGEX plasmids (*Lamin* plasmid was provided by Kristen Johansen). *E. coli* BL21 competent cells were transformed with the plasmids, grown to an OD600 of ∼1.0 and induced with 0.1 mM IPTG for 2 hours. To generate proteins without the GST tag, GST-Lamin and GST-dKeap1 proteins were treated with PreScission Protease (APExBIO). GST pull downs were conducted using Glutathione Sepharose 4B beads (Cytiva 17075601) according to the accompanying protocol. GST-tagged Lamin or dKeap1 proteins were bound to beads and incubated untagged dKeap1 or Lamin, respectively. The GST-protein complexes were eluted from the beads and analyzed by western blotting.

### Bimolecular Fluorescence Complementation (BiFC) assay

*Drosophila* Schneider 2 (S2) cells were cultured according to standard protocols (Thermo Fisher Scientific) and transfected with *pMT-GAL4* (*Drosophila* Genomics Resource Center) and *pUAST* plasmids containing YN and YC fusions using the calcium phosphate transfection kit (Invitrogen). 10 mg of each construct were used. After 24 hours, the cells were spun down, resuspended in PBS, and imaged using a Zeiss LSM 710 confocal microscope.

### Genetic Assays

To generate *Lamin* overexpression flies in combination with or without *dKeap1* knockdown, *UAS-GFP-Lamin* were crossed with either *tub-GAL4/TM6,Tb,Sb,Hu,e,Dfd-YFP* or *dKeap1*^*EY5*^,*tub-GAL4/TM6,Tb,Sb,Hu,e,Dfd-YFP*. L1 larvae were cultured on apple juice plates and appropriate genotypes (*UAS-GFP-Lamin/+; EY5, tub-GAL4/+* and *UAS-GFP-Lamin/+; tub-GAL4/+*) were sorted using a Leica MZ10 F fluorescence stereomicroscope based on the ubiquitin *GFP-Lamin* fluorescence and the absence of the *Dfd-YFP* marker. Larvae were counted daily to assess the number of larvae that survived to specific stages. T-tests were used for statistical analysis.

## Funding

This work was supported by the National Institutes of Health grant (GM128143) to H.D.

## Author contributions

H.D. and J.C. designed the project and planned the experiments. J.C., E.N., and I.C. conducted the experiments. J.C. and H.D. interpreted the data and wrote the paper.

## Acknowledgments

We thank Dirk Bohmann for generously providing *Drosophila* stocks. We thank Kristen Johansen for generously providing *Lamin* constructs.

## Competing interest statement

The authors declare that they have no competing interest.

---

**Expanded View Figure 2.**
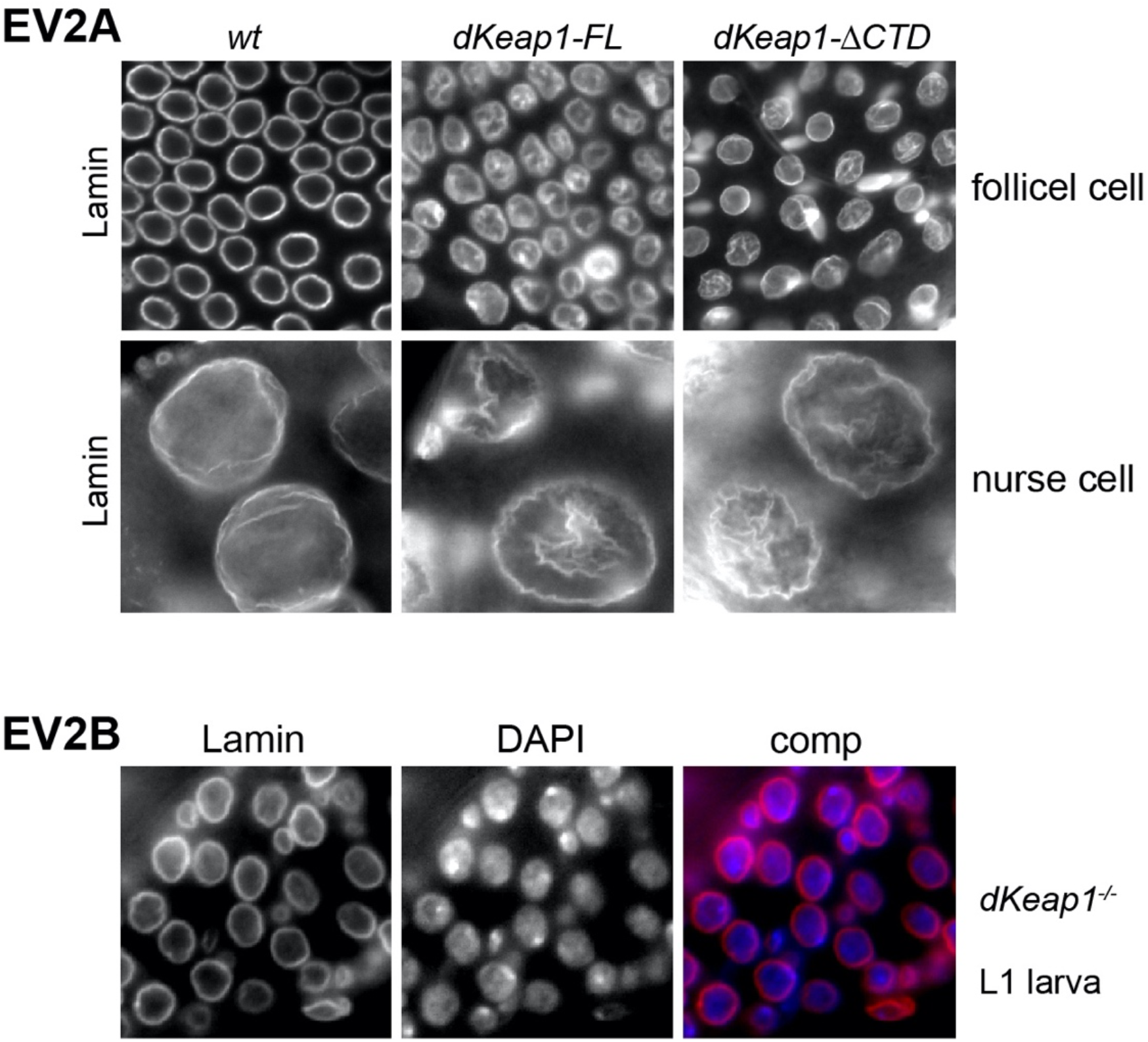

**Expanded View Figure 3.**
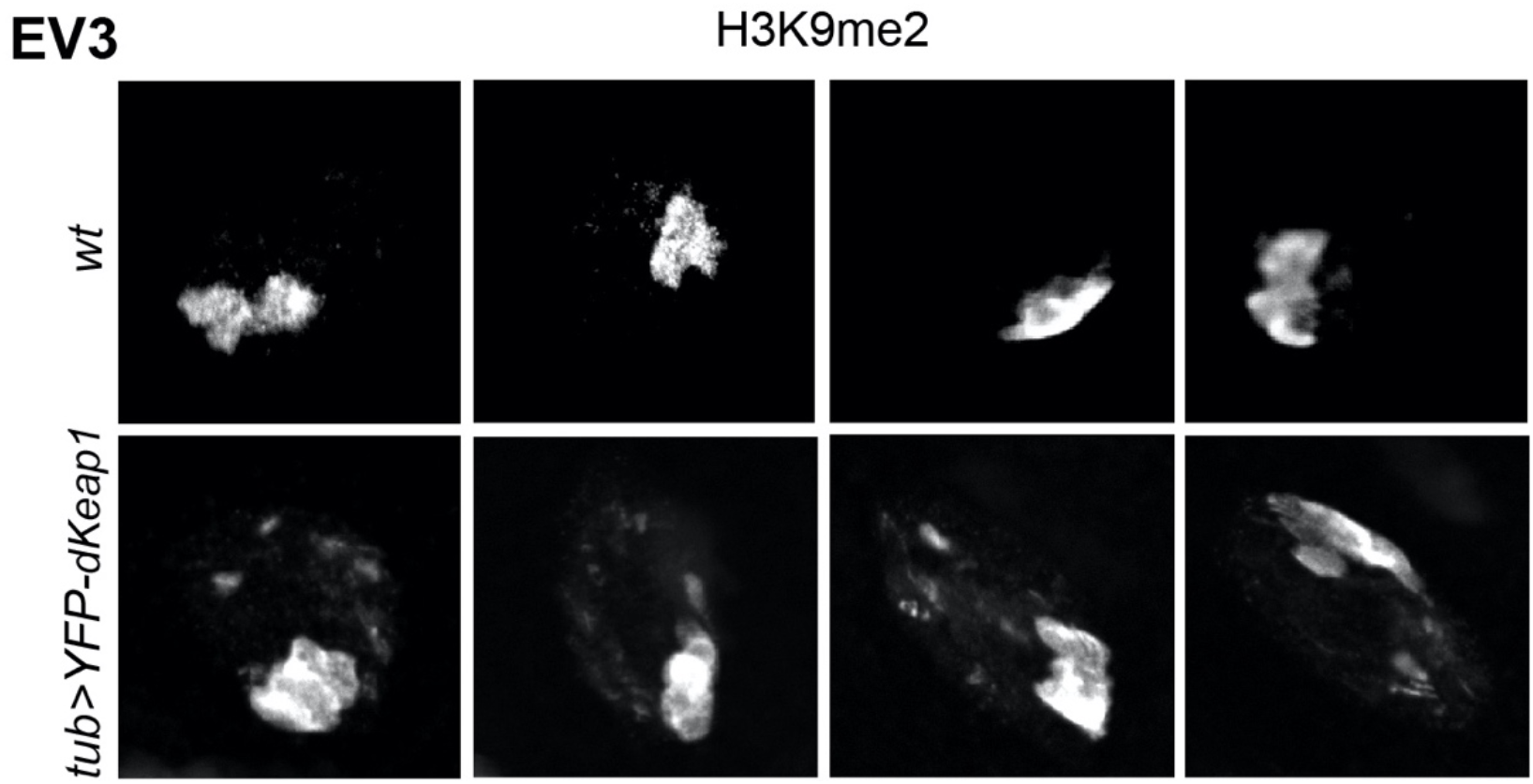

